# QTL mapping for aggressiveness variation in *F. graminearum* revealed one causal mutation in FgVe1 velvet protein

**DOI:** 10.1101/2020.06.19.161349

**Authors:** B. Laurent, M. Moinard, C. Spataro, S. Chéreau, E. Zehraoui, R. Blanc, P. Lasserre, N. Ponts, M. Foulongne-Oriol

## Abstract

*Fusarium graminearum* is one of the most frequent causal agents of the *Fusarium* Head Blight in the world, a cereal disease spread throughout the world, reducing grain production and quality. *F. graminearum* isolates are genetically and phenotypically highly diverse. Notably, remarkable variations of aggressiveness between isolates have been observed, which could reflect an adaptive potential of this pathogen. In this study, we aimed to characterize the genetic basis of aggressiveness variation observed in an F1 population, for which genome sequences of both parental strains are available. Aggressiveness was assessed by a panel of *in planta* and *in vitro* proxies during two phenotyping trials including, among others, disease severity and mycotoxin accumulation in wheat spike. One major and single QTL was mapped for all the traits measured, on chromosome I, that explained up to 90% of the variance for disease severity. The confidence interval at the QTL spanned 1.2 Mb and contained 428 genes on the reference genome. Of these, four candidates were selected based on the postulate that a non-synonymous mutation affecting protein function may be responsible for phenotypic differences. Finally, a mutation was identified and functionally validated in the gene *FgVe1*, coding for a velvet protein known to be involved in pathogenicity and secondary metabolism production in several fungi.

## Introduction

*Fusarium* Head Blight (FHB) is a devastating disease affecting cereals, and is considered to be a limiting factor for global production (Stack, 2000). In addition to yield losses due to fungal infection, grains are often contaminated with stable fungal secondary metabolites toxic for eukaryotic cells, also called mycotoxins, which presence threatens consumers’ health. The hemibiotroph ascomycete *Fusarium graminearum* is the most frequently encountered agent of FHB in the world (Trail, 2009; Van der Lee *et al.*, 2015). *F. graminearum* reproduces through both clonal reproduction and sexual outcrosses, and spores can be dispersed from short to long-range scales (Osborne and Stein, 2007), creating frequent new haplotypes that can widely and rapidly spread under favorable conditions (McDonald and Linde, 2002). Hence, populations of this pathogen show important levels of diversity and gene flow within populations (Akinsanmi *et al.*, 2006; Schmale Iii *et al.*, 2006; Gale *et al.*, 2007; Talas *et al.*, 2012; Prussin *et al.*, 2013; Keller, Bergstrom and Shields, 2014).

The complex interactions between *F. graminearum* and its cereal hosts involve several genetic factors from both partners, the outcomes being quantitative resistances by cereals (Buerstmayr *et al.* 2009) and aggressiveness - defined herein as the quantitative variation of the pathogenicity - by *F. graminearum* (Gilbert, Abramson, McCallum, *et al.*, 2002; Talas *et al.*, 2012; Fabre *et al.*, 2019). This interaction is dynamic, and emergence of more aggressive *F. graminearum* populations has been observed in the fields (Ward *et al.*, 2008; Foroud *et al.*, 2012; Kelly and Ward, 2018). It suggests adaptation followed by a spread of some isolates in their environment, including adaptation to FHB resistant breeds. A detailed description of aggressiveness and its genetic basis, not to mention other part of *F. graminearum* life cycle as dissemination or overwinter survival, is therefore necessary to understand and anticipate its evolution and its consequences (Anderson *et al.*, 2010; Lannou, 2012).

As a whole, every life-related trait during the life cycle of *F. graminearum* that will modulate its level of pathogenicity over time participates in its aggressiveness (Pariaud *et al.*, 2009). Hence, aggressiveness is hard to fully estimate, and proxies like disease severity measurement are often used to assess it. For example, several studies have highlighted important variations of disease severity between and within field isolates (Gilbert *et al.*, 2002; Cumagun and Miedaner, 2003; Talas *et al.*, 2012). For some fungi, mycotoxins can be important aggressiveness component, by killing plant cells and helping their conversion into resources for fungal growth (Pariaud *et al.*, 2009). *F. graminearum* strains are able to produce mycotoxins of the type B trichothecene family (TCTB), either deoxynivalenol (DON) and derivatives (3- and 15-acetylated, 3-ADON and 15-ADON respectively) or nivalenol and derivatives depending on the strain chemotype (Lee *et al.*, 2002; Alexander *et al.*, 2011; Varga *et al.*, 2015). Some TCTB are considered to be aggressiveness factors since their nature and quantities produced has been associated to symptoms severity variation (Bai *et al.*, 2002; Cumagun, 2004; Foroud *et al.*, 2012). But this relationship is not always straightforward, and some correlation is not always observed (Gilbert *et al.*, 2002; Maier *et al.*, 2006; Fabre *et al.*, 2019). In addition to toxin biosynthetic gene clusters, gapless whole-genome sequencing of the four chromosomes of the species associated with extensive genome-mining efforts have highlighted a large repertoire of genes that could be involved in pathogenicity (Cuomo *et al.*, 2007; Brown *et al.*, 2012; Sieber *et al.*, 2014; King *et al.*, 2015). Re-sequencing analyses highlighted high levels of intra-species polymorphism on these *loci*, further suggesting that putative pathogenicity genes are easily affected by mutations, which could play a role in aggressiveness variation and fungal adaptation (Walkowiak *et al.*, 2015; Laurent *et al.*, 2017; Kelly and Ward, 2018).

Several genetic *loci* in *F. graminearum*’s genome have been associated to aggressiveness variation. For example, 50 single nucleotide polymorphisms (SNPs), affecting 26 different genes sparsely located on the genomes of 119 German field isolates have been associated with disease severity variation and suggest a complex genetic control (Talas *et al.*, 2016). The possible roles of each of these genes in aggressiveness has, however, not been functionally validated yet. Moreover, none of those described alleles were detected in any of the isolates originating from France that we have previously characterized for their contrasted levels of deoxynivalenol production in wheat causing symptoms of different intensities on spikes (Pinson-Gadais *et al.*, 2013; Laurent *et al.*, 2017). Therefore, we hypothesized that the genetic basis of aggressiveness variation between these latter strains was different. To test this hypothesis, we developed a QTL-mapping approach using two previously sequenced strains to produce a recombinant population, for which several phenotypic traits related to aggressiveness were assessed. QTL-mapping aims to identify the genetic factors based on their position on the genome, estimate their effect and their heredity, and was complemented in this study with genomic and functional approaches to identify the causal mutation(s) underlying the QTL position(s).

## Results

### Parental strain description

A full-sib family was constructed after crossing the strain INRA-171 and the strain INRA-156 for which the gene *mat-1-2-1* was deleted to bypass homothallism restriction, and named hereafter as INRA-156Δ*Mat* (see Experimental procedure). Deletion of *mat-1-2-1* gene has no effect on disease severity and TCTB production on wheat (Figure 1a and 1b). Both parental strains were able to infect wheat but INRA-156Δ*Mat* – as well as the wild-type strain-induce significantly more severe symptoms than INRA-171 (Figure 1a and Figure 1c). Both strains produced deoxynivalenol (DON) and one derivative, namely 15-acetylated deoxynivalenol (15-ADON). TCTB concentration in wheat flower tended to be greater for INRA-156Δ*Mat* than for INRA-171 (Figure 1b, *p*-value=0.026). Other contrasted phenotypes were observed on the basis of morphological aspects when grown *in vitro* condition. Thereby, INRA-156Δ*Mat* exhibited fast growing and reddish colored mycelia, referred later as morphotype 1, while INRA-171 showed slow-growing, denser and differently pigmented mycelia, referred later as morphotype 2 (Figure 1d). Morphological differences were also observed under microscope: INRA-156Δ*Mat* showed long and unbranched hyphae, and INRA-171 showed hyper-branched and dense hyphae (not shown). According to these observations and in order to assess the genetic basis of the variation in aggressiveness between these two strains, the progeny was phenotyped during two trials for disease severity on wheat, TCTB production in wheat spike, as well as TCTB production, radial growth, fungal biomass and conidia production from *in vitro* culture.

**Figure 1:**
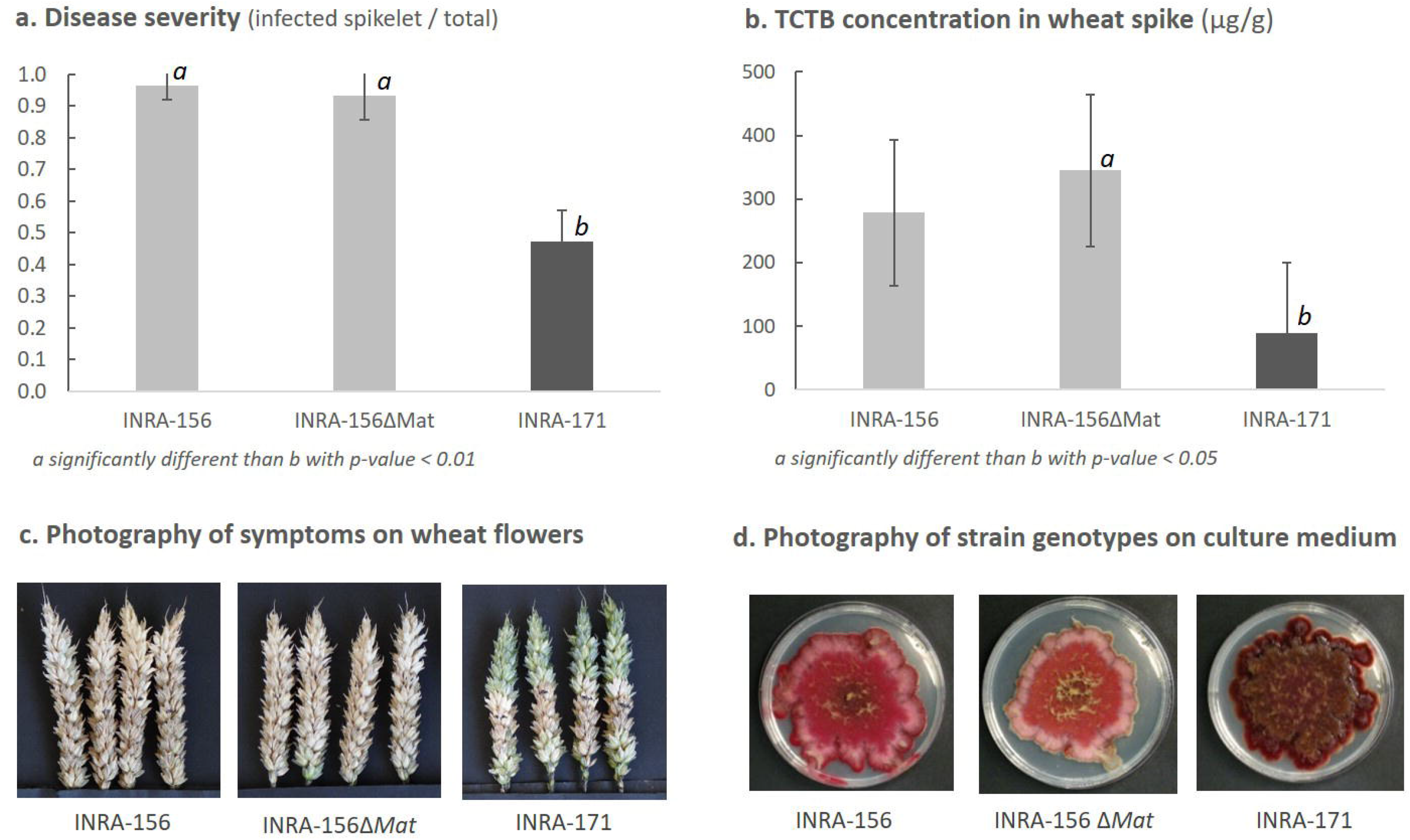
Phenotypic characteristics of INRA-156, INRA-156Δ*Mat* (parental strain), and INRA-171 (parental strain). a. Disease severity (infected spikelet / total spikelet number) measured at 22 days post inoculation (dpi). b. Trichothecene of type B quantification in μg of mycotoxin per gram of dried spike. c. Pictures of Royssac variety wheat spikes inoculated with INRA-171, INRA-156 or INRA-156Δ*Mat* strains. d. Pictures of the strains grown of PDA medium.

### Progeny phenotyping

For each trait measured, the phenotypic values were found always significantly correlated between the two trials, except for the ergosterol measurement (Supplemental File 1A). For clarity purposes, only the analysis of the first trial (T1) will be presented in details herein (Figure 2) and the results of the second trial (T2) are presented in Supplemental file 1. During first trial, the ratio of disease severity varied quantitatively and ranged from 0.3 to 1.0. TCTB concentrations in spike could vary from 27.6 to 493.8 μg per g of dried matter (mainly corresponding to deoxynivalenol and little amount of 15-acetylated derivative) (Figure 2). The normality of the trait distribution was true for the biomass production, the ergosterol concentration and the conidia production (Figure 2, Supplemental file 1A). The distribution of the other traits suggested a bimodal-like distribution (Figure 2, Supplemental file 1A). Interestingly, significant transgressive phenotypes were identified for *in planta* TCTB production (T2 only), disease severity, and both *in vitro* TCTB and biomass production, as well as radial growth (Figure 2, Supplemental file 1A). The morphotypes 1 and 2 were easily distinguished in the progeny and followed a Mendelian 1:1 inheritance (*p*-value <0.01).

**Figure 2:**
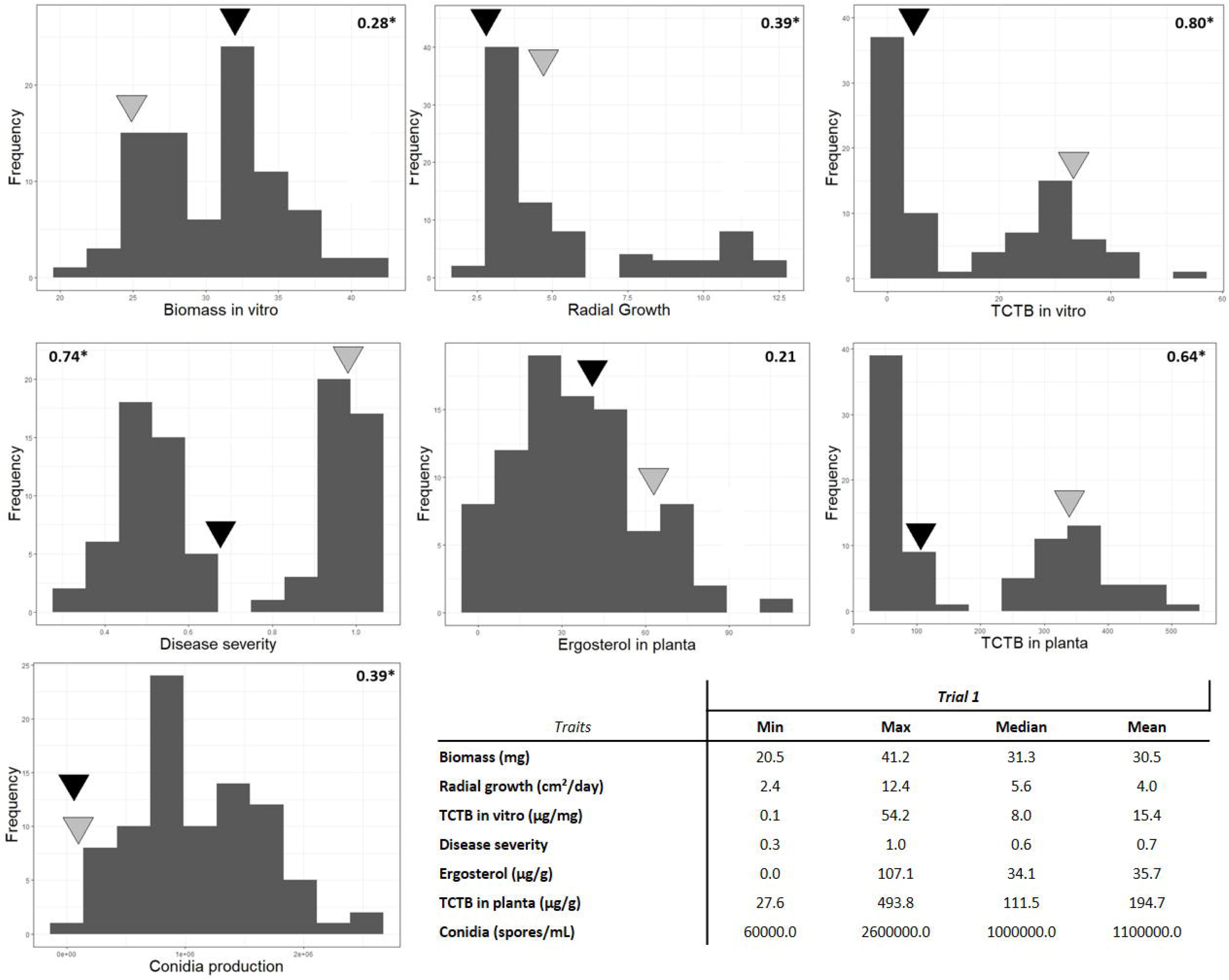
Basic statistics of the traits in the progeny and distribution of the phenotypic traits assessed on the offspring during the first trial. Spearman rank order correlation coefficient between the two trials is in bold for each traits, a star indicates *p*-value < 0.01. Black triangle and grey triangle show the INRA-171 and INRA-156Δ*Mat* values for the trait respectively.

### Principal components analysis and correlation between traits

PCA analysis of the first trial data revealed a strong effect of the first dimension that explains 64.1% of the inertia (Figure 3). The comparison of the inertia explained by this axis with the inertia obtained by the 0.95-quantile of random distributions (22.63%) suggest that only this first axis is informative. As observed on the PCA graph of variables (Figure 3a), disease severity was negatively correlated with biomass production *in vitro* (rho= − 0.66, Supplemental file 1A) but strongly and positively correlated with TCTB production *in planta* (rho=0.78, Supplemental file 1A), TCTB production in *vitro* (rho=0.76, Supplemental file 1A) and with radial growth *in vitro* (rho=0.62, *p*-value < 0.01, Supplemental file 1A). Disease severity was more moderately positively correlated with ergosterol (rho = 0.46, Supplemental file 1A) and conidia production (rho=0.52, Supplemental file 1A). This axis also delineates clearly two group of strains strongly correlated to the morphotypes 1 and 2 (Figure 3b, Wilks’ Lambda test *p*-value 2.56 e-45). Strains of the first group (morphotype 2) typically exhibited low values for symptoms severity, ergosterol, TCTB production *in planta* and *in vitro*, radial growth and conidia production and high values for *in vitro* biomass (Figure 3). The second group of strains (morphotype 1) typically exhibited an opposite situation: high values of disease severity, TCTB *in planta* and *in vitro* and low values for biomass *in vitro* culture (Figure 3). Some strains of this group had high values of ergosterol *in planta* while others had high radial growth and important conidia production. The PCA analysis using the second trial gave similar results (Supplemental file 1B) for the first dimension, although explaining a lower percentage of total inertia (45.3%). This dimension was again highly correlated with the morphotypes (Wilks’ Lambda test *p*-value 6.20 e-32, Supplemental file 1B). A second dimension was considered this time, explaining 19.3% of the inertia (Supplemental file 1B), which opposed genotypes producing high amount of ergosterol *in planta* and high conidia production *in vitro* to individuals with low values of biomass and of conidia production both *in vitro* (Supplemental file 1B).

**Figure 3:**
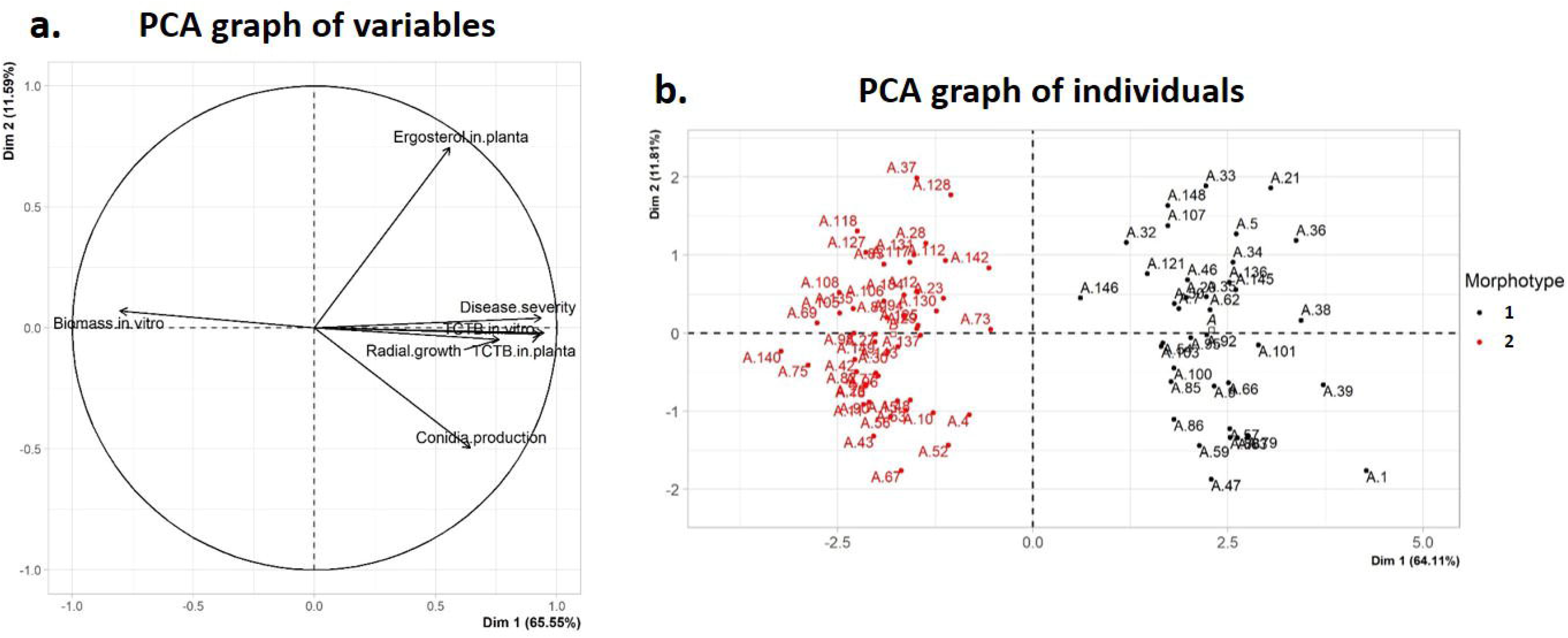
PCA graphs of variables (a.) and of individuals (b.) of the first phenotyping trial data.

### Analysis of variance and heritability estimate

ANOVA from the combined data of the two experiments revealed highly significant genotype effects for all the traits (Table 1). The models also revealed a significant experiment effect and interactions by genotype effect (G × E, Table 1). Considering the correlations observed between the two trials for most of the traits, we considered the experiment effect was more likely due to changes in trait magnitude between the two experiments rather than inconsistent behavior of genotypes. Disease severity and TCTB production *in vitro* showed the best heritability (92.5% and 79.4% respectively, Table 1). The lowest heritabilities were observed for ergosterol concentration (23.0%) and biomass production *in vitro* (31.9%, Table 1). Except for these two traits, the strong heritabilities suggested that the genotype effect was the predominant source of variation for the values of the traits. We focused on the identification of the genetic factors underlying the phenotypic variation through QTL detection. However, given the significant experiment effect, we treated QTL detection by separate experiments for all the traits (Table 2, Supplemental file 1C).

**Table 1:** Analyses of variance and broad sense heritability estimates (h^2^) for the traits measured in planta and in vitro from combined experiment data (two experiments per trait).

| Traits | Disease<br>severity (%) | | TCTB *in planta*<br>($\mu\text{g/g}$) | | Ergosterol<br>($\mu\text{g/g}$) | | Biomass<br>(mg) | | TCTB *in vitro*<br>($\mu\text{g/mg}$) | | Radial<br>growing<br>($\text{cm}^2/\text{day}$) | |
| --- | --- | --- | --- | --- | --- | --- | --- | --- | --- | --- | --- | --- |
| Source of<br>variation | df | MS<sup>a</sup> | df | MS<sup>a</sup> | df | MS<sup>a</sup> | df | MS<sup>a</sup> | df | MS<sup>a</sup> | df | MS<sup>a</sup> |
| G | 93 | 0.29** | 93 | 280123<sup>*</sup> | 93 | 5045** | 93 | 52.4<sup>*</sup> | 93 | 3250** | 93 | 28.5<sup>*</sup> |
| E | 1 | 0.42** | 1 | 1403745** | 1 | 107119<sup>*</sup> | 1 | 34.3<sup>*</sup> | 1 | 36751<sup>*</sup> | 1 | 230** |
| G x E | 92 | 0.02<sup>*</sup> | 88 | 139951<sup>*</sup> | 88 | 3837** | 87 | 31.1<sup>*</sup> | 86 | 668** | 84 | 10.9<sup>*</sup> |
| Residuals | 187 | 0.01 | 183 | 84148 | 183 | 1011 | 364 | 2.8 | 362 | 35 | 339 | 1.2 |
| **$h^2$** | **92.5%** |  | **49.3%** |  | **23.0%** |  | **31.9%** |  | **79.4%** |  | **61.7%** |  |

a: mean square
\*\*: $p$ -value < 0.001
\*: $p$ -value < 0.01

**Table 2:** QTL associated with phenotypic variation under experiment condition and mapping of the morphological Mendelian trait.

| Trait | Chrom | QTL position (cM) | Confidence interval (cM) | Flanking position 5' <sup>a</sup> | Flanking position 3' <sup>a</sup> | LOD score | Thresh-old <sup>b</sup> | Trait variation due to QTL (%) | Favorable allele <sup>c</sup> |
| --- | --- | --- | --- | --- | --- | --- | --- | --- | --- |
| Biomass | I | 90.9 | 89.8 - 92.0 | 3,446,465 | 4,952,659 | 16.0 | 6.1 | 54.2 | B |
| TCTB *in vitro* | I | 92.1 | 90.9 - 93.2 | 4,032,446 | 5,267,806 | 39.0 | 4.9 | 86.5 | A |
| Radial Growth | I | 93.2 | 92.0 - 93.2 | 4,952,659 | 5,267,806 | 10.8 | 5.4 | 43.3 | A |
| Conidia production | I | 92.1 | 90.9 - 93.2 | 4,032,446 | 5,267,806 | 6.11 | 5.4 | 30.7 | A |
| Disease severity | I | 92.1 | 90.9 - 93.2 | 4,032,446 | 5,267,806 | 41.6 | 4.8 | 88.3 | A |
| TCTB *in planta* | I | 92.1 | 90.9 - 93.2 | 4,032,446 | 5,267,806 | 48.0 | 5.0 | 91.5 | A |
| Ergosterol | I | 93.2 | 92.0 – 98.7 | 4,952,659 | 5,368,315 | 6.2 | 5.2 | 26.4 | A |
| Morphological aspect | I | 92.1 | 90.9 - 93.2 | 4,032,446 | 5,267,806 | 25.5 | 4.3 | - | - |

<sup>a</sup> corresponding to the position of the reference genome (RRES v4.0) of the closest marker from one and the other side of the confidence interval
<sup>b</sup> Calculated using 500 permutations
<sup>c</sup> "A" corresponds to INRA-156Δ*mat* strain and "B" to INRA-171 strain, favorable allele increases trait values

### QTL mapping

A single and highly significant QTL that co-localized in the same chromosome region was detected for each trait from each trial, with strong effect for most of the trait (Table 2, Supplemental file 1C). In the first trial, 88.3% of the observed disease severity variation was explained by one single QTL (Logarithm of Odds/LOD= 41.6), located at 92.1cM on the linkage group I corresponding to the first chromosome (Table 2). Similarly, a QTL co-localizing in the same region explained 91.5% (LOD = 48) and 86.5% (LOD = 39.0) of the TCTB production *in planta* and *in vitro* respectively (Table 2). The most moderate QTL effects in trait variation were observed for ergosterol production and radial growth (26.4% / LOD=6.2 and 43.3% / LOD = 10.8 respectively, Table 4). The parental allele increasing the trait values (*i.e.* favorable allele) was brought by INRA-156 strain for all of the traits, except for the biomass measurement *in vitro,* of which the favorable allele was brought by INRA-171 strain (Table 2). These results are consistent between the two phenotyping trials (Supplemental file 1C) and no other significant QTL could be identified with composite interval mapping or with multiple QTL mapping. The morphotype locus was co-localized at the QTL position (Table 2), consistently with the segregation pattern observed for the traits and the results obtained from the PCA analysis.

### *In silico* identification of the causal mutation(s)

The closest markers flanking the confidence intervals highlighted above (90.9 cM – 93.2 cM, Table 2) were aligned on the reference genome (RRES v4.0), delineating a genomic region from the 4,032,446 bp position to the 5,267,806 bp position of the chromosome I of the reference genome (RRES v4.0, Table 2). This region contains 428 putative protein coding genes (Supplemental file 2 for details about the genes). In order to reduce the number of genes, we took advantage of the sequencing data available for the two parental strains and assumed that a non-synonymous mutation affecting the protein function was possibly responsible of the phenotypic differences (Figure 4). In a first step, we identified 67 genes showing allelic polymorphism between the two parents (Figure 4; Supplemental file 2B) of which 46 encoded for polymorphic proteins (Step 2; Supplemental file 2B). Eleven of those were predicted to have altered protein activity due to the presence of the mutation (Step 3; Supplemental file 2B). Of these, 3 were discarded as they shared additional deleterious mutation(s) in both parental strains and were predicted to be not functional in both genotypes. Four additional genes were discarded because the newly described alleles were also identified in strains described previously (Laurent *et al.*, 2017) but showing different phenotypes than the one expected from the parental phenotype. Finally, four candidate genes remained, namely FGRRES_01323 (FGRAMPH1_01G03271), FGRRES_11955 (FGRAMPH1_01G03545), FGRRES_01454 (FGRAMPH1_01G03575) and FGRRES_01553 (FGRAMPH1_01G03787). All have been previously described to be expressed in wheat inoculation condition (Harris *et al.*, 2016) but only FGRRES_01323 and FGRRES_11955 have a predicted biological function (Figure 4, Supplemental file 2B). FGRRES_11955 (FGRAMPH1_01G03545), the orthologue of the *VeA* gene of *Aspergillus nidulans*, was of the highest priority because it is predicted to encode for a major transcription factor, involved in the pathogenicity of several fungal pathogens including *F. graminearum* (Jiang *et al.*, 2011; Merhej *et al.*, 2012). The mutation in *FgVe1* affected the INRA-171 strain and was located on a conserved Velvet domain by mutating the serine located at position 207 into asparagine (Figure 5).

**Figure 4:**
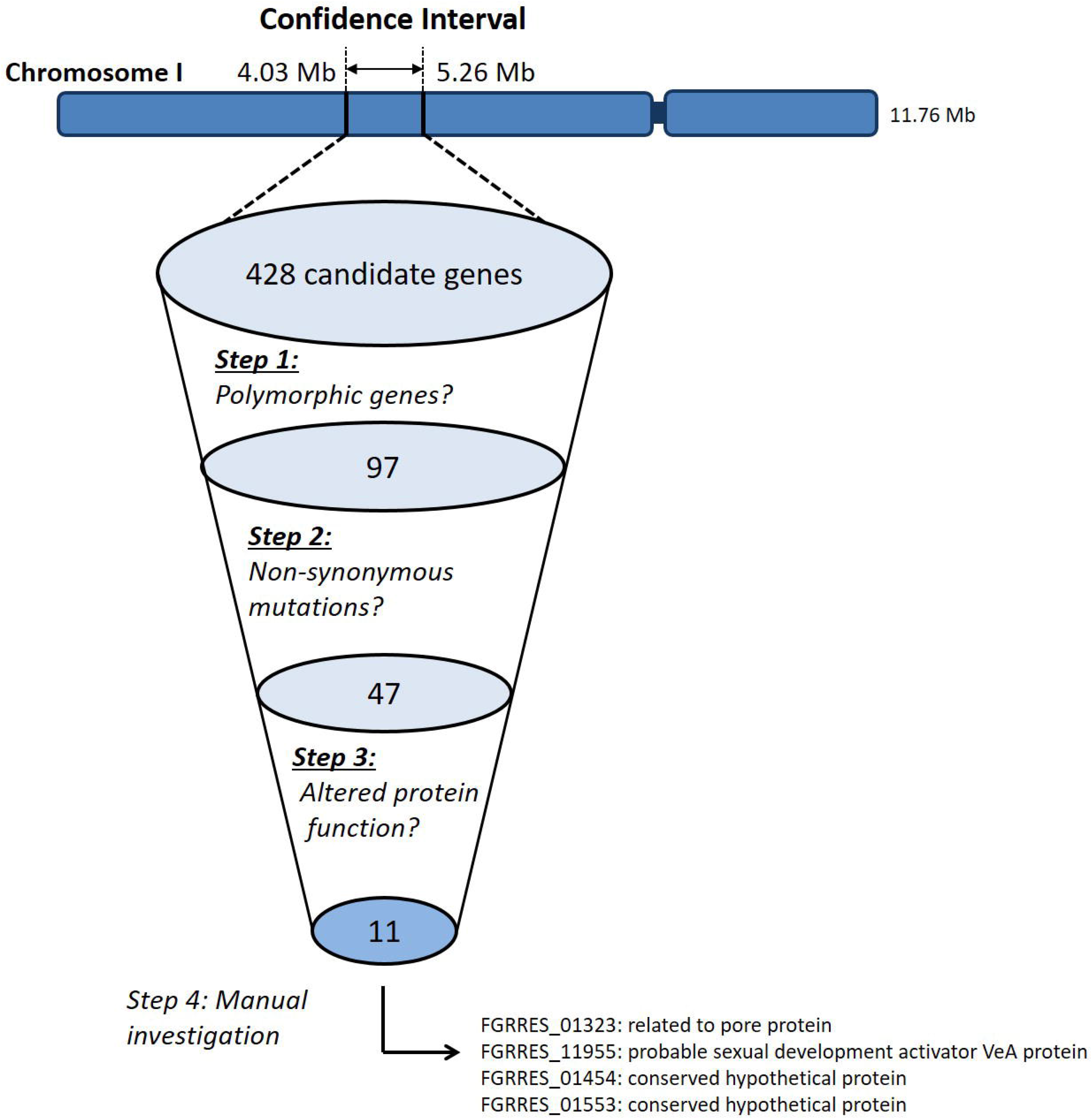
Analytical process of the *in silico* analysis of the genomic region of interest.

**Figure 5:**
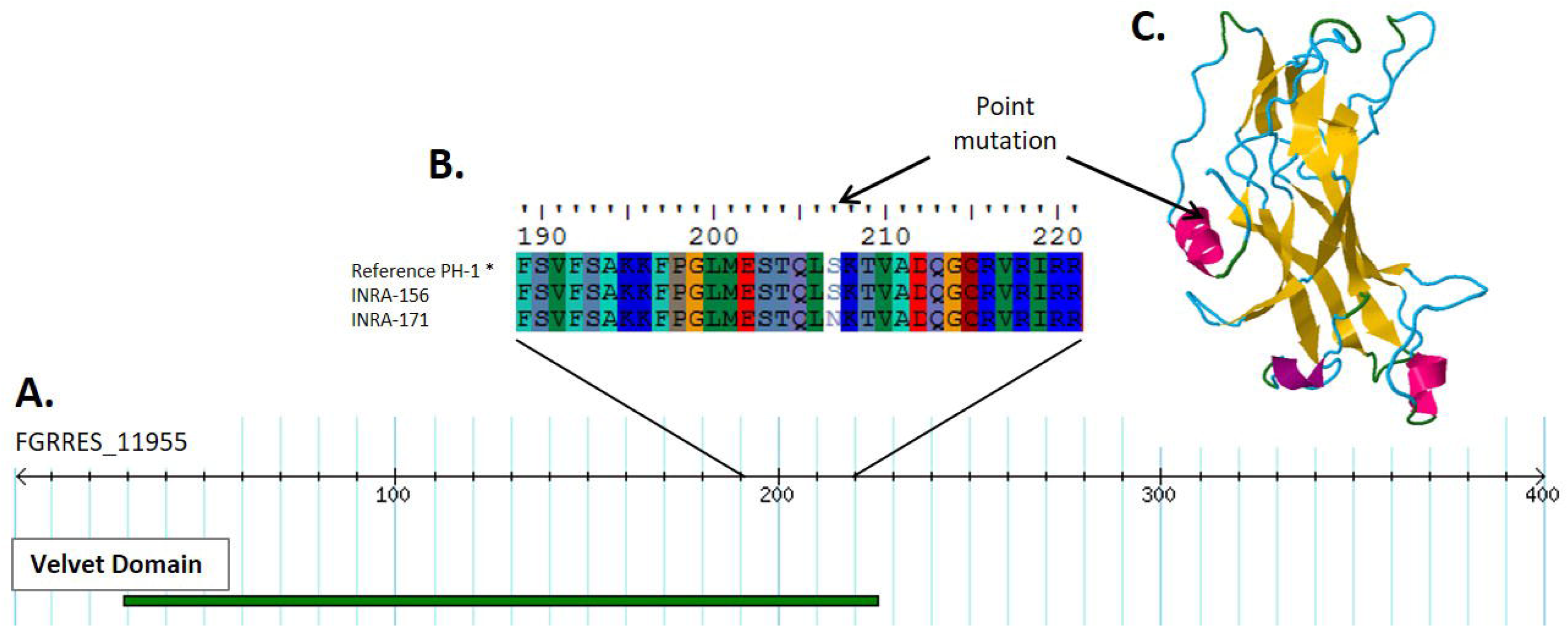
Identification of the causal mutation. A. Representation of the FgVe1 (FGRRES_11955) protein sequence (in amino-acids number) and location and length of the predicted “Velvet domain” (Accession PS51821, Interpro ID: IPR037525) in green. B. Alignment of the protein sequence at the mutation position for the reference genome sequence (* PH-1 strain), for the INRA-156Δ*Mat* and the INRA-171 strain. C. Three-dimensional prediction of the FgVe1 protein and location of the point mutation (dark arrow) on the alpha-loop domain.

### Functional validation of FgVe1

The INRA-156 allelic version of this gene was replaced into the genetic background of the INRA-171 strain. Fungal biomass of the mutant was significantly reduced compared to the wild type strain, as well as significantly reduced compared to the INRA-156 (Figure 6a). The amount of TCTB production for INRA-171*cFgVe1* was significantly greater than INRA-171 strain and INRA-156 strain (tested *in vitro* condition, Figure 6b). Disease severity of the INRA-171 *cFgVe1* was also significantly greater than the wild type and comparable to the INRA 156 strain. INRA-171*cFgVe1* exhibited lighter and more aerial mycelia than the wild-type as observed in the INRA-156 strain (Figure 6d). Overall these results confirm that the mutation identified in the INRA-171 strain was involved in the altered phenotypes.

**Figure 6:**
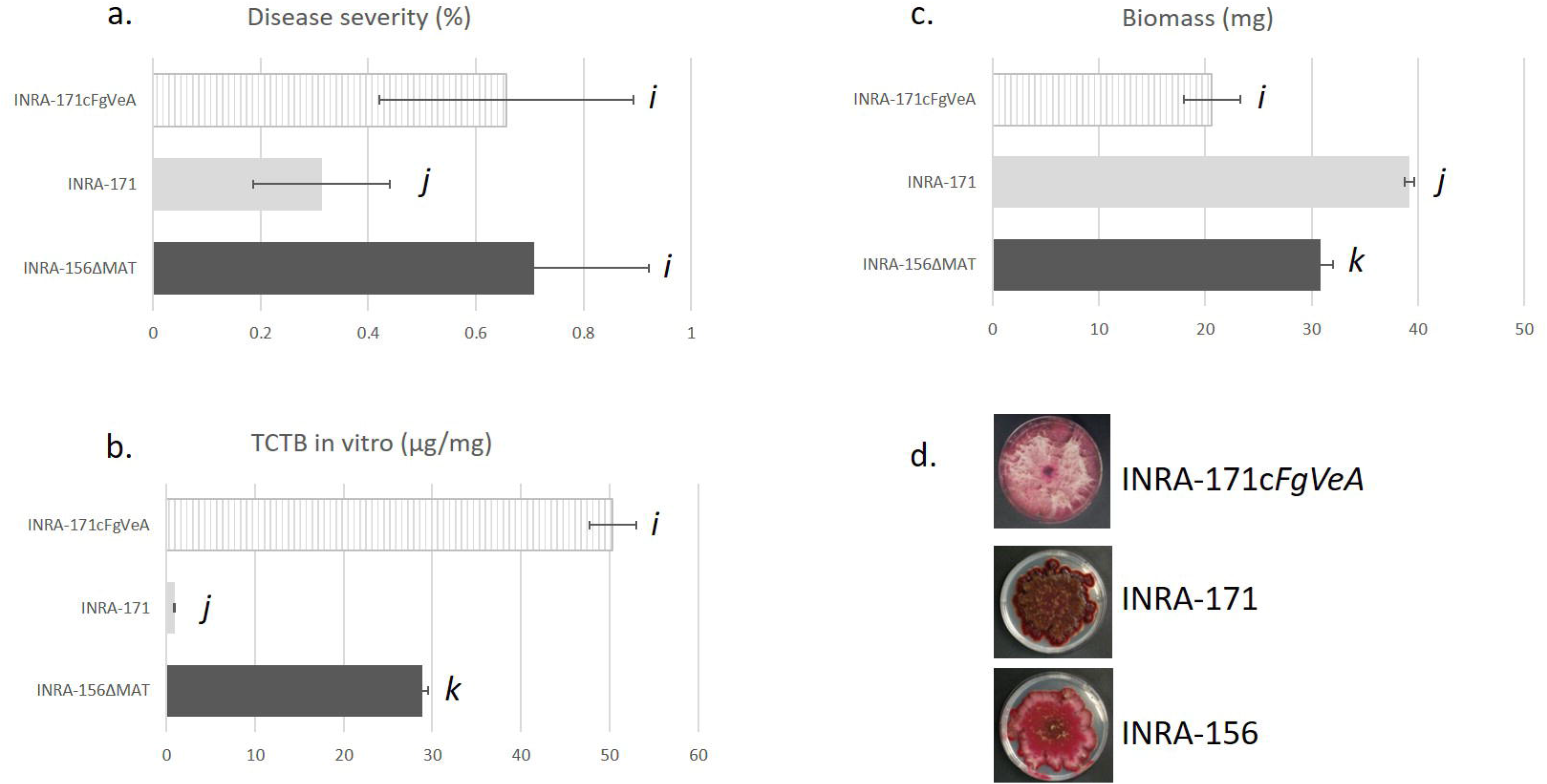
Phenotyping data of INRA-156 strain, INRA-171 strain and INRA-171cFgVe1 strain, of which the *FgVe1* allele has been replaced by the allelic version of INRA-156 strain. a. TCTB production measured from *in vitro* cultures. b. Conidia production from *in vitro* cultures. c. Fungal biomass production from *in vitro* culture. The letters “i”, “j”, and “k” indicates significant differences between the means. d. Pictures of strains grown on PDA medium.

## Discussion

### Different proxies are needed to study aggressiveness

Disease severity on spike is the most commonly used proxy to characterize aggressiveness variation of *F. graminearum* (Cumagun *et al.*, 2004; Talas *et al.*, 2012; Talas *et al.*, 2016). Quantification of deoxynivalenol is also relevant, firstly because of the health issues due to its presence in food and feed, and secondly for its suspected role for plant pathogenicity. During our analysis, we revealed a remarkable correlation between deoxynivalenol concentration and disease severity, as previously reported (Paul *et al.*, 2005; Buerstmayr and Lemmens, 2015). By avoiding the thickening of the plant cell wall and allowing the spread of the fungus (Bai *et al.*, 2002; Jansen *et al.*, 2005; Brown *et al.*, 2010; Walter *et al.*, 2010), deoxynivalenol production is likely involved in a greater colonization of the fungus within the spike. Furthermore, the strong correlation of deoxynivalenol production *in planta* and *in vitro* culture suggested that the strains used in this study exhibited different intrinsic abilities of mycotoxin production, independently of the perception of plant defense. As a whole, the ability of strains to cause symptoms on wheat was clearly related to other intrinsic differences between strains like morphology, radial growth or biomass accumulation. The fact that strains belonging to the morphotypes 2 – that caused little symptoms on wheat - had a reduced radial growth but accumulated the largest biomass *in vitro* was intriguing and suggested different abilities to colonize the plant tissue compared to the morphotype 1. This hypothesis was validated by the measure of ergosterol in wheat spikes, one of the most common proxy used to assess the fungal (Gessner and Schmitt, 1996; Gilbert, Abramson *et al.*, 2002), which showed that strains of the morphotype 1 had a greater spike colonization potential. Despite our attempt to measure conidia production *in vitro* to assess its role for *F. graminearum* aggressiveness, inconsistency between trials were observed, probably due to a strong environmental effect. Finally, others phenotypic traits linked to spore dispersal and germination (including ascospores), infection efficiency, competition with other strains or species, overwinter survival, host or climate adaptation, are necessary to have a more exhaustive picture of the epidemiological potential of fungal pathogens (Montarry *et al.*, 2007; Pariaud *et al.*, 2009; Lannou, 2012; Suffert *et al.*, 2015; Suffert *et al.*, 2018; Desprez-Loustau *et al.*, 2019). Hence, using genetic and genomic approach to investigate the genetic basis of other life-history traits could be addressed in the future to better understand and anticipate the epidemiology of *F. graminearum*.

### Aggressiveness variation is due to a major QTL with pleiotropic effects

In our study, one major effect QTL on chromosome I was detected for each trait that was supported by the bimodal-like distribution observed for the majority of the traits in the progeny (Kang, 2002). This QTL could explain up to 91% of the variation, as for disease severity and TCTB production. Previously, one major QTL explaining 50% of the aggressiveness variation observed in an interspecific cross was associated to the *tri* genes cluster, responsible of the strain chemotype determinism (deoxynivalenol vs nivalenol production), located on chromosome II (Cumagun, 2004). In 2012, one SNP (chromosome I) explaining 26% of the genotypic variance for aggressiveness observed in European field populations was associated to the *MetAP1* gene of unknown function, as well as another SNP associated to *Erf2* (Chromosome IV) - a component of the RAS protein subcellular localization pathway in yeast - explaining 13.1% of the variance. One other SNP (chromosome II) explaining 4.4% of the genotypic variance for deoxynivalenol production in the same populations was associated to the *Tri1* gene, a P450 oxygenase previously characterized and involved in the deoxynivalenol biosynthesis (McCormick *et al.*, 2004). In 2016, 50 and 29 quantitative trait nucleotides were identified in German field populations to be implicated in aggressiveness and DON production variation respectively (Talas *et al.*, 2016). Maximum effect of QTN was 24% for aggressiveness and 19% for DON production and no pleiotropic QTN for these two traits was detected (Talas *et al.*, 2016). The positions of the QTL identified in the present study differ from these previously characterized *loci*, supporting the existence of diverse mechanisms underlying the aggressiveness of *F. graminearum*. Furthermore, the QTL identified herein affects several traits related to aggressiveness, including disease severity, TCTB production, growth and morphology. Such QTL, with pleiotropic effect, are frequent in the nature (Mackay *et al.*, 2009). This result is consistent with the ability of *F. graminearum* to rewire rapidly its physiology thanks to its tightly interconnected and complex network of gene regulation (Lysoe *et al.*, 2011; Guo *et al.*, 2016; Brown *et al.*, 2017; Guo *et al.*, 2020). The presence of this major QTL does not exclude the existence of additional QTL with minor effects that could not have been detected in our experiment. Increasing the population size, or use strains from other genetic backgrounds, would possibly reveal additional QTL at other positions (Mackay *et al.*, 2009).

### The identification of FgVe1 causal mutation highlights the added value of the next generation sequencing in QTL mapping approach

Firstly, next-generation sequencing approaches increase the power of mapping strategies, by facilitating high-throughput genotyping and leading to high-resolution linkage maps. Secondly, it opens great perspectives to identify candidate genes by giving access to the full parental sequences and study the implication of polymorphisms in genes (Grünwald, *et al.*, 2016; Plissonneau *et al.*, 2017). For example, candidate genes involved in the pathogenicity of *A. nidulans* towards human were identified on the basis of QTL location and the presence of non-synonymous mutations within genes (Christians *et al.*, 2011). A similar approach was developed in *Zymoseptoria tritici* to identify candidate genes implicated in aggressiveness in wheat (Stewart *et al.*, 2016), melanin accumulation (Lendenmann *et al.*, 2014), thermal adaptation (Lendenmann *et al.*, 2016), fungicide sensitivity (Lendenmann *et al.*, 2015), or oxydative stress resistance(Zhong *et al.*,2020), and for which some candidate genes were later identified (Krishnan *et al.*, 2018; Meile *et al.*, 2018). Even though QTL mapping strategy remains rarely use to study the genetic basis of pathogenicity of fungi, these examples illustrate their relevancy (Plissonneau *et al.*, 2017). The QTL detected in this analysis was located in a genomic region previously reported to be non-recombinant (Cuomo *et al.*, 2007; Laurent *et al.*, 2018). In consequence to the lack of recombination events in the zone, the confidence interval of the QTL was large (~1.2Mb) and the number of genes (428) was too large to consider functional studies. In order to encompass this limitation, the QTL detection was embedded by genomic and bioinformatics predictions. By doing so, *FgVe1* was identified as the highest-priority candidate, highlighting the power of this mixed strategy to go deeper into the molecular mechanisms underlying trait variation.

### The Velvet proteins are a well known master regulator

VeA proteins are conserved master regulators in several fungi and act in complex with the VelB protein to form the Velvet complex (O. Bayram *et al.*, 2008; Calvo *et al.*, 2016). In *A. nidulans*, the velvet complex was shown to coordinate fungal development, fruiting body formation and secondary metabolism in response to light through the regulation of histone methylation mediated by LaeA (Ö. Bayram *et al.*, 2008). In *F. graminearum*, *FgVe1* has been previously shown to affect fungal development, asexual reproduction, lipid metabolism and secondary metabolism, including trichothecenes biosynthesis (Jiang *et al.*, 2011; Merhej *et al.*, 2012). Therefore, mutant strains deleted for this gene were affected for pigmentation, showed a reduced hydrophobicity and aerial mycelium production, an important decrease of DON production and became non-pathogenic (Jiang *et al.*, 2011; Merhej *et al.*, 2012). The spontaneous mutation affecting *FgVe1* in this study replaced one serine by an asparagine in a conserved α-loop, previously suggested to bind DNA (Ahmed *et al.*, 2013). This mutation was predicted to have no effect on the three-dimensional structure of the protein itself. In regards to the similitudes of the phenotypes observed in INRA-171 with previous mutants (Jiang *et al.*, 2011; Merhej *et al.*, 2012), we hypothesized that the DNA-binding site specificity or activity in this newly described *FgVe1* gene allele is altered. Nonetheless, the aggressiveness of the strain with such allele is reduced, and not totally suppressed. The natural mutant identified herein is a rare opportunity to investigate more deeply the function of Velvet proteins in *F. graminearum* and more widely in fungi.

### What can we say about FgVeA polymorphism?

The polymorphism of the FgVe1 locus was never identified previously neither from the previous genome-wide analysis (GWAs) conducted in Germany (Talas *et al.*, 2012, 2016), nor under selection in natural North America populations (Kelly and Ward, 2018). Data from French populations are still missing, but the screening of a larger French collection of *F. graminearum* (Pinson-Gadais *et al.*, 2013) of about a hundred of srains isolated mainly from diseased wheat, failed to identify this mutation in other genetic background (data not shown). This allele could have been created during strain passing in laboratory (King *et al.*, 2015) or naturally created and maintained at rare frequency in population of *F. graminearum* isolated during the growing season of cereals. Rare alleles are frequently involved in the phenotype variation of living organism, and sometimes accounting for large effect in trait variation (Maher, 2008; Mackay *et al.*, 2009; Gorlov *et al.*, 2011). Noteworthy, if *FgVe1* polymorphism occurs naturally at rare frequency, its identification would have been challenged by GWA, since allele with minor frequency are often filtered out due to statistical constraints (Gorlov *et al.*, 2008). Hence, QTL detection studies, although investigating a limited source of the diversity of a population, can be a powerful and complementary strategy to GWA to identify alleles underlying complex trait variation. Interestingly, another strain from this French collection, namely INRA-195, was also affected by a missense mutation and shared similar phenotypic characteristics than INRA-171. In that strain, the mutation replaced the glycine at the position 118 by aspartic acid and was predicted to have deleterious effect (Table S3 in Laurent *et al.*, 2017). Preliminary results suggest that this new mutation is responsible of the observed phenotypes. This new observation is intriguing, and question the role of Velvet proteins for *F. graminearum* biology and adaptation, but more investigations are needed at that stage to clarify these points.

## CONCLUSION

By using a complementary strategy of QTL detection and comparative genomic analysis, we were able to identify one causal and non-synonymous mutation affecting the DNA binding site of the FgVe1 protein and leading to major differences of disease severity on spike, DON production, fungal growing and morphological characteristics. The discovery of *FgVe1* polymorphisms rises new interrogations about their origins and its evolutionary consequences, that will be addressed in future researches.

## Experimental procedures

### Parental strains information and progeny construction

A description of the parental strains and of the construction of progeny is described in (Laurent *et al.*, 2018). Briefly, the two parental strains INRA-156Δ*Mat* and INRA-171 have been isolated in France and belong to the DON/15-ADON chemotype. INRA-156Δ*Mat* was constructed from INRA-156 strain of which the gene *mat-1-2-1* (FGRRES_08893) was replaced by a hygromycin resistance cassette. Ninety-four strains from the progeny were used for phenotyping trials among which 88 were used for linkage map construction. Previous inoculation of INRA-156 and INRA-156Δ*Mat* strains was conducted on wheat spikes and showed no significant effects on the disease severity and TCTB production in wheat due to the *mat-1-2-1* deletion (Figure 1). The parental wilt-type strains were previously whole-genome sequenced, as well as 4 other *F. graminearum* strains (Laurent *et al.*, 2017).

### Phenotyping

For each trait assessment, two experiment trials were conducted as time repeats for the entire progeny, referred hereafter to as T1 and T2.

#### Disease severity assessment

Two bread wheat cultivars (*Triticum aestivum*) were used for *in planta* trials. The moderately FHB susceptible cv. Royssac for the *Fusarium* progeny phenotyping, and the FHB susceptible cv. Apogee for the functional validation of *FgVeA* gene. Plants were grown under greenhouse, as previously described (Fabre *et al.*, 2019). Plant duplicates were conducted per progeny strain according to the protocol described above. Three to four spikes per plant showing the same ontogeny were inoculated at mid-anthesis. A conidial suspension of 100 spores adjusted in 10μL of carboxymethylcellulose medium (CMC, Carboxylmethyl cellulose 15%, yeast extract 1%, MgSO4,7H2O 0.5%, NH4NO3 1%, KH2PO4 1%) was deposited in two floral caveats of two opposite spikelets per spike. Plants were inoculated with medium and served as control. The severity ratio corresponded to the number of infected spikelets per spike and normalized by the total number of spikelets per spike, as previously suggested (Talas *et al.*, 2012). This trait was measured at 22 days post inoculation (dpi). Spikes were collected following notation in order to be dried and used later on for TCTB and ergosterol quantification. In addition to these phenotypic traits, the latent period and the area under the disease progress curve were also measured during the first trial and are not shown as they strongly correlated to the disease severity measurement at 22 dpi.

#### In planta TCTB assay

TCTB were extracted from 1g of dried spike powder. Powder was diluted in 5 mL of Milli-Q water and the solution was vortexed for 10 min, and then centrifuged 12 min at 5000rpm. 1mL of this solution was diluted in 3 mL of Milli-Q water. 7.5μL of this solution was suspended in 742.5μL of a solution containing 78% of H20 and 22% of methanol, before filtration through 0.22 μm-pore-size filters (Phenomenex, Torrance, USA). Quantification of trichothecenes was performed using a QTrap 2000 LC-MS/MS system (Applied Biosystems) equipped with a 1100 Series HPLC system (Agilent), as described by (Atanasova-Penichon *et al.*, 2018) with slight modifications. Chromatographic separation was achieved at 45°C with a Kinetex™ 2.6 μm XB-C18 column (150 × 4.6 mm, Phenomenex, Torrance, USA). Solvent A consisted of methanol/water (10/90, v/v) and solvent B consisted of methanol/water (90/10, v/v). The flow rate was kept at 700 μl/min and was split so that 350 μL/min went to the electrospray source. Gradient elution was performed with the following conditions: min with a linear gradient from 85 % to 5 % A, 4 min held at 5 % A, 1 min linear gradient from 5 % to 85 % A and 85 % A for 8 min post run reconditioning. The injection volume was 5 μl. Mass spectrometry conditions were identical to that described by Atanasova-Penichon *et al.* (2018).

#### In planta ergosterol assay

extraction and quantification of ergosterol was performed according from (Touati-Hattab *et al.*, 2016): with small modifications: dried samples were suspended in 100 μl of methanol and filtered through a 0.2 μm filter (Phenomenex, Torrance, USA); quantification was performed on a Shimadzu Prominence UFLC chain. Ergosterol was measured in μg/g of spike powder.

*In vitro TCTB assay* conidial suspension of 125,000 spores (0.1 - 0.6mL) were incubated in a final volume of 8mL completed with Minimum Synthetic (MS) medium (previously described by (Boutigny et al., 2009) for 14 days at 25°C, in dark conditions. Triplicates of cultures were conducted per strain. The cultures of liquid media were transferred in tubes (15 mL) and centrifuged at 5000rpm at 4°C. Supernatants were retrieved and 250μL per sample were collected, diluted in 250 μL of MeOH and filtered using 0.22 μm-pore-size filters (Phenomenex, Torrance, USA). Quantification of TCTB was performed on a Shimadzu Prominence UFLC chain as described by (Hadjout *et al.*, 2017)

#### Radial Growing

mycelium was grown on Potatoes Dextrose Agar (PDA 39g/L, Difco-France) at 25°C in dark condition. This phenotyping assay was conducted using automated image analysis procedure. Briefly, strains were grown on triplicate from size-standardized mycelia plugs, and pictures were taken every day using standardized camera settings and lighting environments. Images were analyzed using ImageJ software (Schneider *et al.*, 2012). Pictures were stacked and length were scaled manually using 5cm tracks disposed during picture acquisition. All pictures were treated simultaneously using the “montage” option and fungal surfaces were automatically selected using saturation and brightness thresholds. The delimitated surfaces, representing fungal radial growing, were integrated using ROI manager and areas were measured in cm². Radial growing was measured in cm² per day.

#### In vitro biomass

fungal biomasses resulted from MS medium cultures and *in vitro* production of mycotoxin assay presenting above. Triplicates were conducted for each strains and fungal biomasses were dried during 5 days at 80°C. Biomasses were measured in mg.

#### Conidia production

4 plugs of mycelium grown previously on PDA medium (~ 0.5cm diameters each) were incubated in 12mL of CMC liquid medium for three days at 25°C and under permanent shake (180rpm). Singleton of conidial suspensions was produced. After incubation, conidial suspensions were filtered through Sefar Nitex 03-100 (100 μm, SEFAR AG - Switzerland). Filtered solutions were measured for optical density at 500nm. Quantification was performed by external calibration using conidial solutions of which concentrations of spores were counted using Thoma cell.

### Statistical analysis prior and during QTL detection

Statistical difference between the mean values of the parental phenotypes was tested using ANOVA and the post-hoc multiple comparisons of means using the Tukey’s Honestly Significant Difference method with the Multcomp package in R. Correlations between experiments and between traits have been tested using Spearman rank order test and considered as significant for *p*-value < 0.01. Shapiro Wilk test was used to test the normality of the distribution of the trait in the progeny and considered as normal for *p*-value > 0.01. PCA analysis were conducted using FactoMineR and Factoshiny packages in R version 3.6.1 (R Core Team; 2020). Mendelian inheritance was tested using Chi-square test and was accepted for *p*-value < 0.01. Statistical difference between the mean values of the progeny phenotypes and the parental strain ones was tested using ANOVA and the post-hoc multiple comparisons of means using the Dunnett’s method and the Multcomp package in R. Phenotypes significantly greater or lower than the parental ranges (*p*-value < 0.01) were considered as transgressive. Broad sense heritabilities were estimated considering combined data according to the model: *Y* = *μ* + *G* + *E* + *G* × *E* + *ε*, with G, the genotypic effect, E, the experiment effect, *G* × *E* the interaction between genotypic effect and experiment effect and *ε* the residual effect. The linkage map used for QTL mapping was previously described in Laurent, *et al.*, 2017. QTL detection was conducted using R/qtl (Broman *et al.*, 2003) and the interval mapping analysis in R version 3.6.1. The Haley-Knott regression method was set (Haley and Knott, 1992). Missing marker genotyping data were filled with the imputation method proposed by Rqtl, using the Kosambi map function and an estimate genotyping error rate of 0.001. QTL were accepted if the logarithm of the odds (LOD) score at the position was greater than the LOD threshold calculated with 500 permutations and for a significance level α of 0.001. Confidence intervals corresponded to the +1/−1 positions (in cM) of the maximal LOD score of the QTL, expanded to the closest markers at the outer edge of this interval. Markers were aligned on the reference genome of *F. graminearum* RRES v4.0 (King *et al.*, 2015).

### *In silico* analysis of candidate genes

Genomic sequences and genetic variants of parental strains were retrieved from Laurent *et al.*, 2017. In order to reduce the number of candidate genes, an analytical process was designed and summarized in Figure 4. This process proposes to identify polymorphic genes (Stage 1), affected by non-synonymous mutations (Stage 2). Variant annotation was conducted using SnpEff software (Cingolani *et al.*, 2012), as previously described (Laurent *et al.*, 2017). To predict if an amino-acid substitution or InDels have an impact on the biological function of the protein (Stage 3), Provean algorithm was used (Choi *et al.*, 2012) and polymorphic proteins with a score equal or less than −2 were kept as candidate proteins with highly probable altered activity. In Stage 4, the remaining candidates were filtered out on the basis of the presence of additional deleterious mutation found for both parental strains and compared to the genotypic information from strains, previously characterized (Laurent *et al.*, 2017). Hence, genes having common candidate mutation(s) with other strains should have comparable phenotypes, *e.g.* if the candidate gene has one mutation in the INRA-171 parent (inducing symptoms of little severity on plant and producing low level of TCTB) and was identify in a strain with a highly aggressive and toxinogenic, this candidate mutation was filtered out (Supplemental file 2B). Gene information and orthologues from other species were retrieved from the FungiDB database (Stajich *et al.*, 2012).

### *FgVe1* validation and investigation

*FgVe1* locus from INRA-156 was first amplified using the P1 primer (AAGAAACATCTCCGCCATTTTA) and the P2 primer (TGAAACAAAAATTCTGCCAATG). Amplicon length was 3.4 kb and corresponded to *FgVe1* coding and flanking sequences (FGRRES_11955). Kapa Hifi™ DNA polymerase (KapaBiosystems) was used for amplification, following these conditions: 95°C 2 min, 35 cycles of [98°C 20 s, 61.8°C 15 s, 72°C 1 min 45 s], 72°C 5 min. PCR product was then purified using the Wizard® SV Gel and PCR Clean-up System (Promega) and quantified using NanoDrop™ technology. The PCR product was mixed with plasmid pBC HygromycinR to co-transform protoplasts of *F. graminearum* strain INRA-171. Protocol for protoplast production and transformation was used according to Montibus *et al.*, 2013. Transformed strains were selected on hygromycin B containing PDA medium (60 μg/mL) and single germinating conidia were isolated and transferred on new PDA plates. Integration of the INRA-156 allelic version of the gene *FgVe1* instead of the INRA-171 allele, and removal of the INRA-171 allele, was verified by Kompetitive Allele Specific PCR markers genotyping (KASP™, Supplemental experimental procedure). The genetic background of the transformed strains was verified using 4 additional KASP markers distributed on other chromosomes (Supplemental experimental procedure). Phenotypes of the transformed strains were tested as previously described in “Phenotyping” section. Two- and three-dimensional structures of the Velvet proteins were predicted from the corresponding amino-acid sequence of the two allelic versions of Velvet gene using PhyrT2 (Kelley *et al.*, 2015). Protein structures were visualized using Jmol (Hanson *et al.*, 2010). Difference between the mean values of the phenotypes between the parental strain and the mutant strain was tested using ANOVA and post-hoc multiple comparisons of means using Tukey’s Honestly Significant Difference method and significance was accepted for *p*-value < 0.01.

## Supporting information

A. Phenotypic statistics and correlation between trials and traits. B. Distribution of the different phenotypes assayed in the offspring during the se

Result from the analytical process proposed in Figure 4 and aiming to identify the candidate gene underlying the QTL. A) Genes located in the confiden

Phenotypic and genotypic data formatted for R/qtl analysis

## Acknowledgement

BL received a PhD fellowship from the French Research Ministry. Authors would like to gratefully thank Cyrille Saintenac for helpful conversations and for assistance during *in planta* experimentations; Ross Houston and Christos Palaiokostas for remarkable advices during work conduction and along manuscript writing; Pierre Chesnot for providing a valuable help during English writing, Cécile Robin for helping reviewing the manuscript. The authors have no conflict of interests to declare.

## Data Availability Statement

The data that support the findings of this study are openly available in https://data.inrae.fr at https://doi.org/10.15454/IRKFHM. Genotypic and phenotypic data are also available in Supplemental file 3. The short-read sequences of the wild type parental strain genomes (INRA-156 and INRA-171) are available at the GenBank Sequence Read Archive under the accession SRP064374.

## Supporting Information legends

**Supplemental file 1:** A. Phenotypic statistics and correlation between trials and traits. B. Distribution of the different phenotypes assayed in the offspring during the second trial. Dark triangle and grey triangle show the INRA-171 and INRA-156Δ*Mat* value for the trait respectively. C. Principal component analysis for the different traits. D. QTL detection for the different traits measured.

**Supplemental file 2:** Result from the analytical process proposed in Figure 4 and aiming to identify the candidate gene underlying the QTL. A) Genes located in the confidence interval of the QTL. B) Annotation of the polymorphism between the two parental stains inside the QTL region.

**Supplemental file 3:** Phenotypic and genotypic data formatted for R/qtl analysis

